# Bistable bacterial growth dynamics in the presence of antimicrobial agents

**DOI:** 10.1101/330035

**Authors:** Nelly Frenkel, Ron Saar Dover, Eve Titon, Yechiel Shai, Vered Rom-Kedar

## Abstract

**Background:** The outcome of a given antibiotic treatment on the growth capacity of bacteria is largely dependent on the initial population size (the Inoculum Effect, IE). For some specific classical antibiotic drugs this phenomenon is well established in both in-vitro and in-vivo studies, and its precise mechanisms, its clinical implications and its mathematical modelling are at the forefront of current research. Traditional view of the IE is that it is mainly attributed to β-lactam antibiotics in relation to β-lactamase producing bacteria, and that some antibiotics do not induce an IE at all. The study of antimicrobial peptides had emerged in the past two decades as a possible additional strategy for combatting infections, and their mechanism of operation and clinical implications are extensively studied. Yet, no previous studies addressed the possible induction of IE under the action of classical cationic antimicrobial peptides (CAMPs).

Based on mathematical reasoning regarding bacteria-neutrophils interaction, we hypothesized that CAMPs also induce an IE in bacterial growth, and questioned what are the similarities and differences between the IE induced by CAMPs and that induced by classical antibiotics. To this aim we also needed to better understand the characteristics of the IE induced by classical antibiotics.

**Principal Findings:** We characterized and built a model of in-vitro IE in *E. coli* cultures using a large variety of antimicrobials, including 6 conventional antibiotics, and for the first time, 4 cationic antimicrobial peptides (CAMPs). Each combination of bacterial initial load and antimicrobial concentration experiment was done in duplicate, with 48 such combinations in each experiment. Each experiment was repeated 4-6 times, sometimes with some adjustments in the tested concentrations to get better resolution of the IE. Each growth curve was processed independently, to correctly reflect the initial exponential growth that might lead to large deviations even between duplicates. By using Optical Density (OD) to monitor the bacterial density, we were able to gather growth curves from this extensive data set and from these curves extract, by data processing, the corresponding growth functions. We show that this process allows us to clearly differentiate between simple one-dimensional deterministic bacterial growth dynamics and more complex behaviour.

In all agents we tested, including all cationic antimicrobial peptides and all conventional antibiotics, independently of their biochemical mechanism of action, an “inoculum effect” was found. At a certain range of concentrations, which is specific for every drug and experimental setting, the system exhibits a bistable behaviour in which large loads survive and small loads are inhibited. Moreover, we characterized three distinct classes of drug-induced bi-stable growth dynamics and demonstrated that in rich medium, CAMPs correspond to the simplest class, bacteriostatic antibiotics to the second class and all other traditional antibiotics to the third, more complex class. In particular, for the first two classes, of cationic antimicrobial peptides and of the commercial bacteriostatic antibiotics, the bacterial growth can be explained by a very simple deterministic one-dimensional mathematical model. These findings provide a unifying universal framework to describe the dynamics of the inoculum effect induced by antimicrobials with inherently different killing mechanisms.

Limitations of the results: The IE we detect is in-vitro, in rich medium, and the simple deterministic one dimensional models apply to this setting for the CAMPs and the bacteriostatic antibiotics only. While these findings can be used as a building block to more complex settings, with in-vivo being the most complex of all, it is clear that additional studies are needed in order to address these complexities. Another limitation is the OD methodology which does not clearly differentiate between live, dormant and dead cells and also does not detect small bacterial loads that are below the reader detection level. Nonetheless, since only live bacteria grow, the growth functions that we find experimentally are independent of the dead and dormant bacteria, and the bacterial density axis may be at most shifted by small amount due to this effect. The behaviour at small loads, below the OD detection level, is also irrelevant for the current study as we are concerned with the IE at high inoculum. Finally, this study is conducted at the population level only, with the point of view that IE is induced by deterministic non-linear interactions between the bacteria and the anti-microbial agent, without delving into the details of the particular molecular mechanisms that lead to this particular interaction. Such detailed nonlinear molecular mechanisms that induce IE are known to exist for some of the agents we use. Future studies are needed to better understand the detailed molecular mechanisms in the other cases.

**Conclusions & Significance:** The vast increase in bacterial resistance, highlights the need for new approaches to eradicate bacterial infections, by either the development of new antimicrobial agents, or new strategies of treatment. Developing treatment strategies requires a better understanding of the Inoculum Effect (IE). We demonstrate that IE is abundant in the application of both classical antimicrobial peptides and classical antibiotics to bacteria. Furthermore, we show that IE falls into three universality classes of bi-stable behaviours and that classical antimicrobial peptides form a class of their own – the simplest and most predictable class. These findings propose a new exciting viewpoint on the universality features of IE that may serve as building blocks for the design of better treatment strategies for infection.

We stress that the detection of IE in CAMPs may have important implications for their mode of operation, and this finding may lead to further explorations of this phenomenon both in terms of mechanistic models and in terms of clinical and biological implications.

While bacterial IE was identified in previous studies of particular conventional antibiotic agents and bacteria, previous explanations of its appearance included genetic and/or phenotypic population heterogeneity and additional time-dependent factors. These were modelled, for example, by deterministic multi-dimensional equations of classical reaction kinetics. Here we show that for some cases (the bacteriostatic antibiotics) a one dimensional model can explain the resulting growth curves by density dependant mechanisms alone. By Ockham’s razor principle, we assert that such models are adequate for describing the IE in bacteriostatic antibiotics. On the other hand, we also show that for all other cases (growth with all other classical antibiotics and growth in poor medium) simple one dimensional deterministic models cannot describe the dynamics, and thus multi-dimensional models may be needed to describe IE in these cases. Additionally, contrary to some other studies, we show that IE appears in every antibiotic we tested (in particular antibiotics that are not β-lactams), so additional molecular mechanisms for creating the non-linear bacterial-drug interaction need to be identified.

Finally, density dependent phenomena are abundant in biology and may appear in other pathogenesis systems, where densities matter. Here we demonstrated that such phenomena can sometimes be described by very simple growth dynamics. Such simple models may serve as building blocks to more complex models such as in-vivo ones and may also inspire detailed studies aimed at deciphering the specific dominant molecular mechanisms of the detected IE. We propose that the principles and methodologies developed here for studying IE by observing the population level dynamics may be applicable to diverse biological situations.

**Authors Summary:** The vast increase in bacterial resistance highlights the need for new approaches to eradicate bacterial infections, by either the development of new antimicrobial agents, or new strategies of treatment. Since the outcome of a given antibiotic treatment on the growth capacity of bacteria is largely dependent on the initial population size (Inoculum Effect, IE), developing treatment strategies requires a better understanding of this effect. We characterized and built a model of this effect in *E. coli* cultures using a large variety of antimicrobials, including conventional antibiotics, and for the first time, cationic antimicrobial peptides (CAMPs). Our results show that all classes of antimicrobial drugs induce an inoculum effect. Moreover, we characterized three distinct classes of drug-induced bi-stable growth dynamics and demonstrated that in rich medium, CAMPs correspond to the simplest class, bacteriostatic antibiotics to the second class and all other traditional antibiotics to the third, more complex class. These findings provide a unifying universal framework to describe the dynamics of the inoculum effect induced by antimicrobials with inherently different killing mechanisms. These findings propose a new exciting viewpoint on the universality features of IE that may serve as building blocks for the design of better treatment strategies for infection.

## Introduction

Many factors can affect bacterial susceptibility to antibiotics. These include the metabolic state and the presence of persistent cells [1–3], the microenvironment conditions that affect the antibiotic potency [4], the physical structure of the population (biofilms) [5], and the population size, or inoculum, at the site of infection. Indeed, a major power of bacteria is within numbers since it has been well established that as a population, bacteria often survive a concentration of an antimicrobial agent that is lethal to individual cells. In a therapeutic context, this means that the fate of an initial infection depends on the initial load of bacteria – while small infections are easily cleared even with no antibiotics, large infections are hazardous, even when antibiotics are administered at high doses. This phenomenon, known as the “inoculum effect” (IE), is well established *in-vitro* [6–8], as well as *in-vivo* in animal models and in human patients [9–11][7, 12] [13] [14]

There are several known biological mechanisms that were proposed to account for IE. First, it is known that the *E. coli* spontaneous beneficial mutation rate is 10^#x2212;5^ mutations per genome per generation [15]. Thus, bacterial populations equal to or larger than 10^5 may contain *genetic heterogeneity* in regard to antibiotic resistance [16, 17]. *Phenotypic heterogeneity* [1–3] is also more prominent in these larger initial populations. It is believed that this heterogeneity can generate bi-stability: at low numbers we expect that only one, non-resistant population exists, hence treatment leads to extinction, whereas at high initial numbers heterogeneity may allow a resistant strain to grow, leading to bi-stability.

Second, *density-dependent mechanisms* may also lead to IE. The bacterial density affects both the cellular state of the cell, and its interactions with the antimicrobial agent. Cellular communication that is sensitive to population density is called quorum sensing. Quorum sensing enables bacteria to synchronize gene expression and alter their properties to become more resistant to different antibiotics. At low cell densities, a large proportion of the signalling factors disperse before they can be used, and so their production provides a small direct or indirect fitness benefit. At high cell densities, a much greater proportion of the signalling factors is available per cell, and consequently bacteria can better cope with the relevant stressor [18]. A density-dependent mechanism that does not involve cellular signalling is reduction in the antibiotic concentration. For instance, *E. coli* secrete the β-lactamase enzyme that cleaves and inactivates β-lactam antibiotics [19]. Larger populations produce more β-lactamase and can therefore degrade the antibiotic faster [20, 21]. If the population survives and grows, more enzymes are produced and cleave the antibiotic in a higher rate - a positive feedback mechanism. A similar concept applies when an antibiotic agent binds to its target or to non-specific cell components and its effective concentration in the medium is reduced. This might not have much effect when the antibiotic concentration is sufficiently large compared to the population size, but is expected to lead to a bistable situation at a critical ratio. Even if the antibiotic concentration does not change in time, bistable behaviour can arise when growth is proportional to the amount of antibiotic molecules available to each bacterium at the time of exposure [7, 22]. The above density dependent mechanisms are inherently non-linear: these effects are not increasing gradually, proportionally to the bacterial load and antimicrobial concentration as they all have either threshold or limiting effects.

In a clinical situation, IE is undesirable because the treatment outcome may become difficult to predict even if the dynamics (drug-target interaction) is assumed to be deterministic [8, 23–26]. A direct consequence is that treatment with insufficient doses of antibiotics can lead to increased mortality of infected patients [27], and favour the selection of drug-resistant strains [25, 28].

The IE may be viewed as a bistable effect. A system is said to be monostable if it always equilibrates to one final state, and is bistable when it admits more than one stable state (see e.g. [29]). In the bacteria-antimicrobial context, a monostable system corresponds to the case where any initial amount of bacteria (*B*) reaches the same maximal population size under a non-lethal treatment (*A*). Lethal treatment leads to eradication of all loads, and therefore, the fate of the system is independent of the initial load (Figure 1A). In a bistable situation, the fate of the system may depend on its initial load. Here, for a range of the antimicrobial agent concentrations, the bacterial population can assume two possible states (the two solid black lines in Figure1B), depending on whether the initial bacterial concentration was above or under the threshold concentration B_c_(A) (dashed black line in Figure1B). Three main regimes govern this bistable dependence. First, a minimal antibiotic/peptide concentration A_c_ is needed to kill or inhibit a minute number of bacterial cells, so for A < A_c_, even minute bacterial populations grow, similar to untreated cells. Second, an enormous amount of antimicrobial agent inhibits practically any amount of cells, so there exists A_e_ such that for A > A_e_ all relevant bacterial densities are growth-inhibited or killed. Third, for A_c_ < A < A_e_, the greater the concentration of the antimicrobial agent, the larger the bacterial concentration it can inhibit, so the threshold concentration B_c_(A) is monotonically increasing with A in this range. It follows that for any given antimicrobial agent concentration in the range A_c_ < A < A_e_, the fate of the bacterial population depends on whether its initial concentration is above or below B_c_(A) (dotted bottom line Figure 1B). We thus have a bi-stable behaviour for all A_c_ < A < A_e_ (Figure 1B).

**Figure 1.**
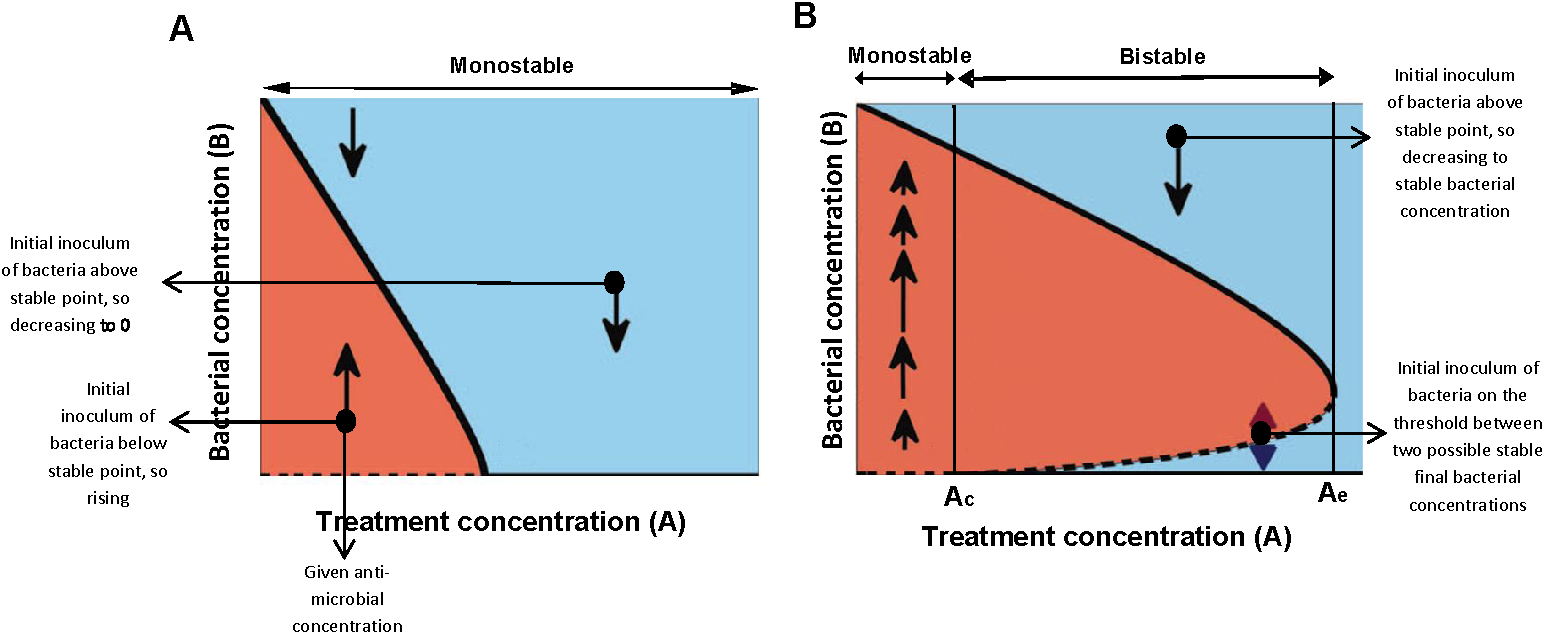
One dimensional models of monostable and bistable bacterial growth in the presence of antimicrobial agents. The x-axis represents the collection of antimicrobial concentrations, the y-axis shows the bacterial amount. In this plot, the bacterial dynamics correspond to motion along vertical lines – from any initial inoculum at a given antimicrobial concentration, the bacterial load increases (moves upward) in the red regions and decreases (moves downward) in the blue regions. The solid/dashed lines correspond to stable/unstable equilibrium states respectively. (A) The monostable model has a single stable equilibrium bacterial concentration for any antimicrobial concentration (a thick full line). (B) The bistable model has a range of concentrations, the bistable range, where the system has at least two possible stable equilibria (two full lines). The upper solid line represents the antimicrobial-concentration dependent maximal capacities. The dashed line of unstable equilibria represents the collection of critical bacterial loads B_c_(A) above which the bacteria grows to the upper equilibrium branch and below which the bacteria becomes extinct or inhibited. Ac is a minimal antimicrobial agent concentration that inhibits/kills a minute number of cells. Ae is a concentration that inhibits/kills every tested bacterial load. The bi-stable range corresponds to concentrations between Ac and Ae. See similar figure in Malka et al [29],[30] for neutrophil-bacteria dynamics.

Essentially, any non-linear drug-target interactions can result in a bi-stable behaviour. Indeed, in Malka et al [29, 30], it was argued, by this reasoning, that neutrophil-bacterial interactions also exhibit bi-stable behaviour. It was shown there that deterministic one-dimensional models, which depend only on a single dynamic variable - the varying bacterial concentration - are sufficient for adequately describing such a behaviour in *in-vitro* experiments in which bacteria and neutrophils are incubated together in well mixed wells. Notably, other more complex mechanisms may also result in a bi-stable behaviour. By Ockham’s razor principle, since the concentration-dependent bi-stable mechanism is the simplest adequate model, and such a model involves hardly any assumptions on the specifics of the neutrophils-bacteria interactions, it provides the main underlying mechanism for *in-vitro* bacteria-neutrophil dynamics (and possibly for *in-vivo* dynamics under neutropenic conditions, see Malka et al. [29, 30]). The inoculum effect shows that a similar underlying mechanism applies to bacteria-antibiotics interactions.

In the current study, we show that as long as the initial bacterial loads used for bacterial growth assays in the presence of antimicrobials were sufficiently resolved, every tested antimicrobial agent showed an inoculum effect for *E.coli* K12 MG1655 cells. We tested several antibiotic substances, each employing a different mechanism to kill bacteria, to show that the bacterial growth dynamics are dependent on the initial bacterial load present in the medium, regardless of the type and mechanism of action of the antimicrobial agent. We also show, for the first time to the best of our knowledge, that classical antimicrobial peptides that do not have specific targets on the bacterial membrane induce an inoculum effect as was previously observed for some non-classical antimicrobial peptides [31]. Moreover, we show that the killing induced by them may be described by an even simpler model – a model of a bistable immediate kill term followed by peptide independent dynamics. Possible clinical implications of this study are discussed.

## Experimental Materials and Methods

### Commercial antibiotics

Ampicillin sodium salt, Kanamycin sulphate, Chloramphenicol, Carbenicillin disodium salt, Oxacillin sodium salt and Gentamycin solution were purchased from Sigma-Aldrich. Polymixin B was purchased from Fluka BioChemika and Tetracycline hydrochloride was purchased from Calbiochem.

### Peptide synthesis and purification

Peptides were synthesized by using a 433A synthesizer (Applied Biosystems, Life Technologies) on rink amide 0.65 mmol/mg MBHA resin, using Fmoc protected amino acids. The synthesized products were washed thoroughly with DMF and DCM, then dried and cleaved. Cleavage was performed by addition of 95% trifluoroacetic acid (TFA), 2.5% water and 2.5% TIS. The peptides were then purified (>98% homogeneity) by reverse phase HPLC (RP-HPLC). Purification was performed using a C18 column and a linear gradient (Melittin – 20%-80%, K6L9 – 10%-90%, MSI – 10%-90%) of acetonitrile in water (final fluid containing 0.1% TFA (v/v)] for 40 min.

### Bacterial Strains

*E. coli* MG1655 with a Lux-Kanamycin resistance plasmid (described previously in [32]) was used for generation of all bacterial growth curves except for the curves generated with Kanamycin where non-resistant *E. coli* MG1655 was used. Bacteria were grown in LB media or LB+Kanamycin (30 µg/ml), shaking in 37°C for 4 hours (OD_600_ = 1-1.5) and then diluted for the experiments.

### Generation of growth curves from various initial inoculums

a sterile 96-well plate with a flat-bottom was prepared with serial dilutions of the necessary antibiotic/peptide in LB media (total volume in well 100 µL). In short, an initial high concentration stock of the antibiotic/peptide in DDW was used to make an initial stock in LB to fill 200 µL into wells in row A. Serial dilutions were then made by a multipipette into the other wells already containing 100 µL of LB (the last row was left without antibiotics/peptide as a positive control). Bacteria from a shaking culture as described above were then measured for their OD_600_ and normalized in 2 ml LB medium to an OD_600_ of 0.01 (order of 106 cfu/ml as measured from live counts). Serial dilutions of the bacteria were then made in additional 5 LB tubes (200 µL from previous dilution were transferred into 1800 µL of LB every time after vortexing). 100 µL of the highest bacterial dilution were inserted into wells A1-B12, 100 µL of the second highest bacterial dilution into wells C1-D12 etc. The smallest inoculum contains about 5 bacteria and, in all experiments` controls, exhibited growth (was not empty). The prepared plate was placed in an automatic microplate plate-reader for 16 hours with medium shaking speed and an OD_600_ measurement every 20 min.

### Antibacterial activity

The base minimal inhibitory concentration (bMIC) was determined for each antibiotic/antimicrobial peptide (AMP) based on the above described growth curves – bacterial populations with an initial inoculum of 10^6^ cfu/ml (105 cfu per well) that finished with an OD_600_ lower than a cut-off of 0.35 after a 16-hour incubation were considered as extinct or inhibited. The lowest antibiotic/peptide concentration for which both duplicates went extinct is the bMIC. Similarly, MICs were also generated for lower inoculums using the same cut-off.

### Preparation of spent media with Polymixin B (PMB) and Ampicillin (Amp)

Appropriate concentrations of PMB and Amp were prepared in LB medium for subsequent dilutions for a MIC assay described above. *E. coli* MG1655 were grown in shaking LB till an OD600 of about 0.4-0.6, and diluted into some of the prepared PMB and Amp stocks to an OD600 of 0.1. All types of these media and LB alone were then incubated in shaking in 37°C for 1h. Incubated media were centrifuged and the supernatant was used for a standard MIC assay described above.

### Determination of antibiotic potency over time - E. coli

MG1655 were grown in shaking LB ON and then centrifuged for 5 minutes in 3000 rpm and diluted to an OD600 of 1. Ampicillin, Chloramphenicol, Carbenicillin and Tetracycline were added to appropriate concentrations for final MIC determination as indicated in the results to tubes containing either LB or LB and bacteria in an OD600 of 1. Some LB tubes with bacteria in the same density were also incubated with rest without any antibiotics. All tubes were incubated for 24 hours in shaking and 37°C. In addition, another inoculum of *E.coli* MG1655 were grown in shaking LB ON. After 23 hours, the whole process was repeated for the 1hr samples with the new ON culture. When 24h had passed, all samples were filtered through 0.2 um syringe filters and appropriate amounts of all corresponding antibiotics were added to the samples that were incubated with bacteria only (without antibiotics). Finally, all samples were diluted by a factor of 2 with fresh LB for the highest antibiotic concentration and by increasing dilutions for a standard MIC assay as described above. In addition, fresh LB with fresh antibiotics was prepared for each antibiotic for a parallel classic MIC assay as a control by the protocol described above. All MIC assays were performed in 96-well plates over 16h of incubation in shaking (250 rpm) and 37°C. Final results were determined by the sight of turbidity and OD600 measurements.

## Mathematical models

Three types of deterministic mathematical models are proposed, all admitting bi-stable behaviour as described in the introduction section: The bistable immediate kill and then A-independent dynamics model (simplest), the bistable A-dependant dynamics model (simple), and a bistable multiple time-dependent factors A-dependant dynamics model (the most complex model presented here). Notably, more complex bi-stable models taking into account detailed molecular processes, stochastic effects, and/or several phenotypic populations may be introduced. One of the main results here is that the two simpler models are adequate for describing much of the data.

Denoted by B(t) (bacterial cells/ml) the concentration of the bacteria at time t (hours) and by A the initial concentration of the antibiotics/peptide (µM). The growth curves under the influence of the antibiotics/peptides reflect the overall population growth which is decreased/delayed by either killing of substantial portions of the population (peptides or bactericidal antibiotics) or by growth-inhibition (bacteriostatic antibiotics).

### The bistable immediate kill model (BIK)

the bacterial destruction occurs quite rapidly and abruptly at some initial phase τ. After this destruction occurs, the surviving bacteria (if such exist) grow exactly as if there are no peptides in the medium (as in the control). The flow chart for such a model is of the form in flow chart No 1.

This means, that if one plots the growth function of the bacteria, dB/dT versus B, one should obtain, for all concentrations of the antimicrobial peptide, a unique growth function F(B) which is independent of the antimicrobial concentration. The experiments (see Figures 3–6) demonstrate that all the antimicrobial peptides exhibit such behaviour whereas none of the antibiotic agents induce such a simple behaviour.

### The bistable A-dependant dynamics model (BAD)

here the bacterial destruction occurs continuously as a result of the antibiotics action, which is assumed to be fixed along the experiment (See flow chart No 2).

The bi-stability arises here from the non-linear nature of antibiotics-bacteria interactions. The experiments (see Figure 3) demonstrate that to a good approximation, two of the antibiotics that were tested are well described by this continuous one dimensional model. From the data sets we can roughly obtain the concentration dependent growth function F(B,A) for each of the antibiotics by plotting the experimental curve 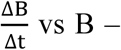 for each concentration separately. Plotting these experimental curves provides a first glimpse to the experimentally derived nonlinear bacterial-antibiotics interactions (see Figure 3B).

### The bistable multiple time-dependent factors A-dependent dynamics model (BMFD)

Here, additional time-dependent factors, such as antibiotics concentrations and β-lactamase concentrations influence the bacterial growth and are being influenced by the bacterial concentrations, leading to *at least* a two *dimensional* deterministic system (it is also plausible that stochastic terms need to be included, see e.g. [26]). The simplest possible model of this kind lets the bacterial dynamics to be modelled as above, with the antibiotics concentration that governs the bacterial dynamics change with time according to a dynamic law – see flow chart No 3.

Clearly, such higher dimensional models are more complex than the deterministic one dimensional BIK and BAD models. Indeed, notice that the two simplified models (BIK and BAD) can be considered as two limiting cases of the above simplest two dimensional model: very fast depletion of the antibiotics leads to a BIK model, whereas very slow depletion and consumption of the antibiotics means that the antibiotic concentration remains essentially fixed and the BAD model emerges.

When the experimental growth function for each antibiotic concentration depends on the bacterial initial load, one concludes that the two simple models are insufficient to describe the bacterial growth dynamics.

**Flow chart No 1.**
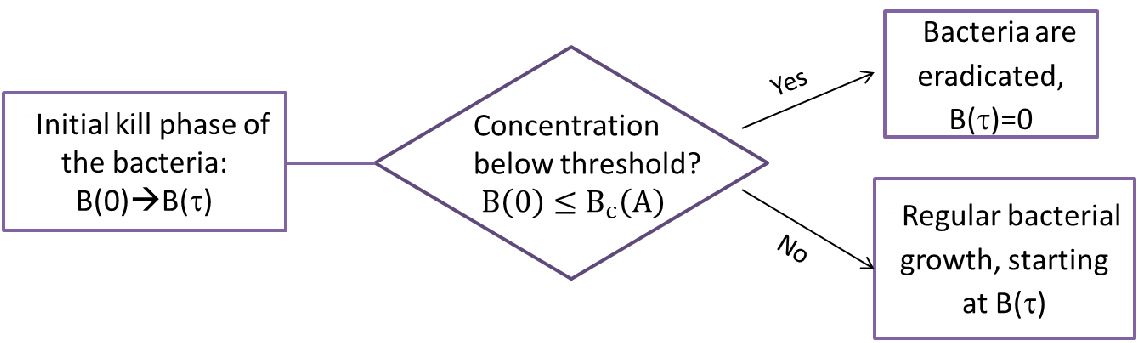
BIK Model.

**Flow chart No 2.**
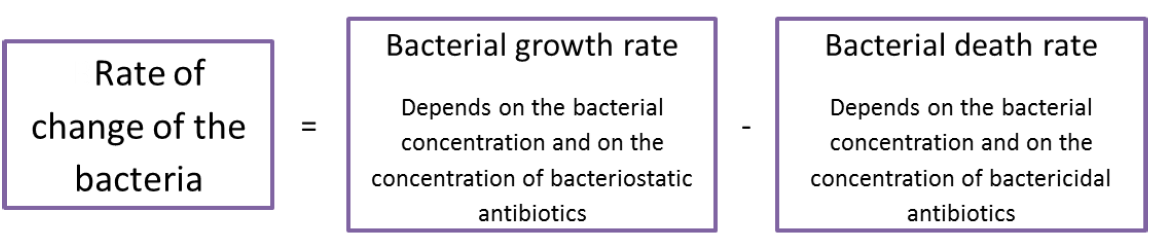
BAD Model.

**Flow chart No 3.**
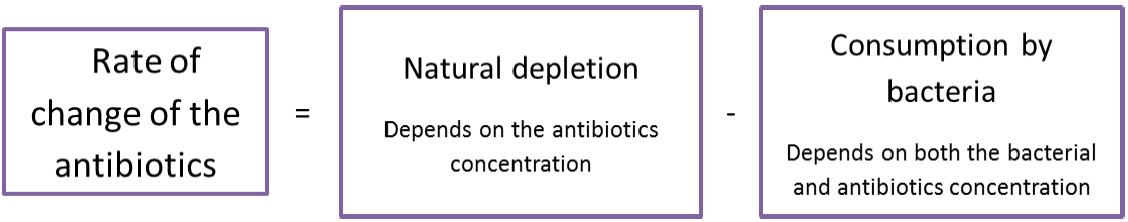
A BMFD Model. Antibiotics dynamics coupled to the BAD model bacterial dynamics

## Results

### Without antibiotics, in rich media, bacteria grow to maximal capacity obeying deterministic 1D growth dynamics

To investigate how the initial bacterial concentration (i.e. initial load) influences the growth outcome at a given antibacterial agent concentration, we first searched for conditions in which the medium content itself is not a limiting factor that can affect the growth dynamics. Six initial bacterial loads with a fold-change between them (roughly 10–10^6 cfu/ml) were prepared from an exponentially growing culture of *E.coli* and grown in a rich medium (LB) or in a minimal medium (see methods). In the rich medium, all initial loads had the same growth rate during the exponential phase, and reached the same maximal concentration (optical density, OD) – the full capacity of the well (Figure 2A). When grown in a minimal medium, the different initial loads did not necessarily reach the same maximal concentration, nor grew with similar rates during the exponential growth phase. In addition, there was a higher variability between replicated samples (Figure 2D). Note that the different loads are detected by the OD reader only after they grow above the detection level of about 10^6-10^7 cfu/ml. The time shift between the different loads can be thus used to estimate the growth rate at the exponential phase (and later on, to estimate the initial anti-microbial killing). To better visualize the specific growth function (dB/dt) for a given bacterial concentration B(t) we plot dOD/dt versus B. In these plots, the growth rates of all loads tested in rich medium converge approximately to a single line (Figure 2B), while in minimal medium they do not (Figure 2D).

**Figure 2.**
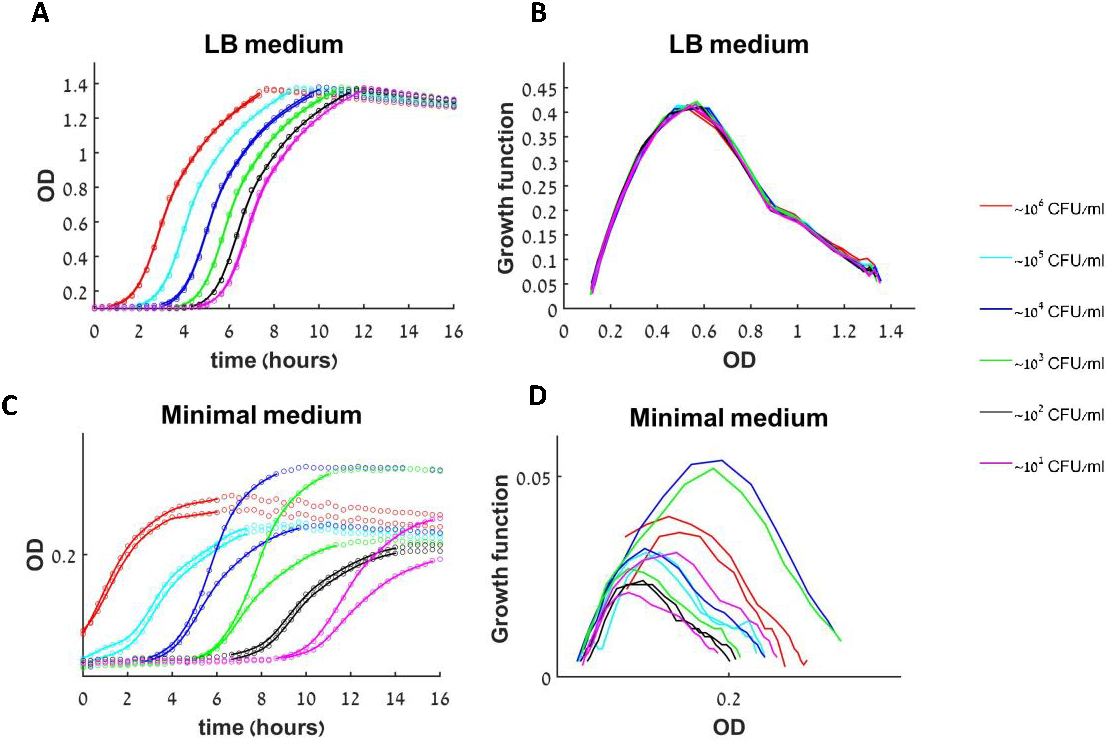
Growth dynamics of untreated bacteria in rich and minimal medium. (**A**) The growth of twelve initial loads (six different loads in duplicated) in rich medium (LB) as monitored by a change in optical density (OD) in time. Circles-data points, solid line – smooth approximation to data used for calculating the growth function, see methods. (**B**) Specific growth function curves [experimental dOD/dt vs. OD]. (**C-D**) Repeating the experiment in minimal medium resulted in higher variability between duplicates and exhibited a strong influence of the initial load on the growth rate and the maximal population size.

These plots show, for the first time (to the best of our knowledge), the fully nonlinear growth function that appears in any deterministic mathematical model which involves bacterial growth. We see that in a rich medium the rate of change of the bacterial concentration (dB/dt) at a given time depends only on its current concentration (B(t)) and not on the initial load B(0) or the time of growth (Fig 2B). Mathematically, such a behaviour corresponds to classical deterministic 1D dynamics, namely to a model of the form dB/dt=F(B) where F(B) is shown in Figure 2B. In particular, the fact that all duplicates and all different initial loads collapse to one growth function clearly demonstrates that under these conditions stochastic effects are insignificant, even when a minute number of live bacteria are put in the wells. In contrast, bacteria in minimal medium exhibit more complicated dynamics – here the rate of change depends on other time dependent factors such as the depleting nutrients, and we observe that duplicates may grow differently (Figure 2D). Thus, the dynamics in minimal medium cannot be described by such a simple mathematical model. We conclude that growth in a rich medium is robust and enables reproducible and identical growth rates for all initial loads, hence we use it to study how antimicrobial peptides and antibiotics with well-defined mechanisms of action (Table 1) affect the growth dynamics.

**Table 1:**
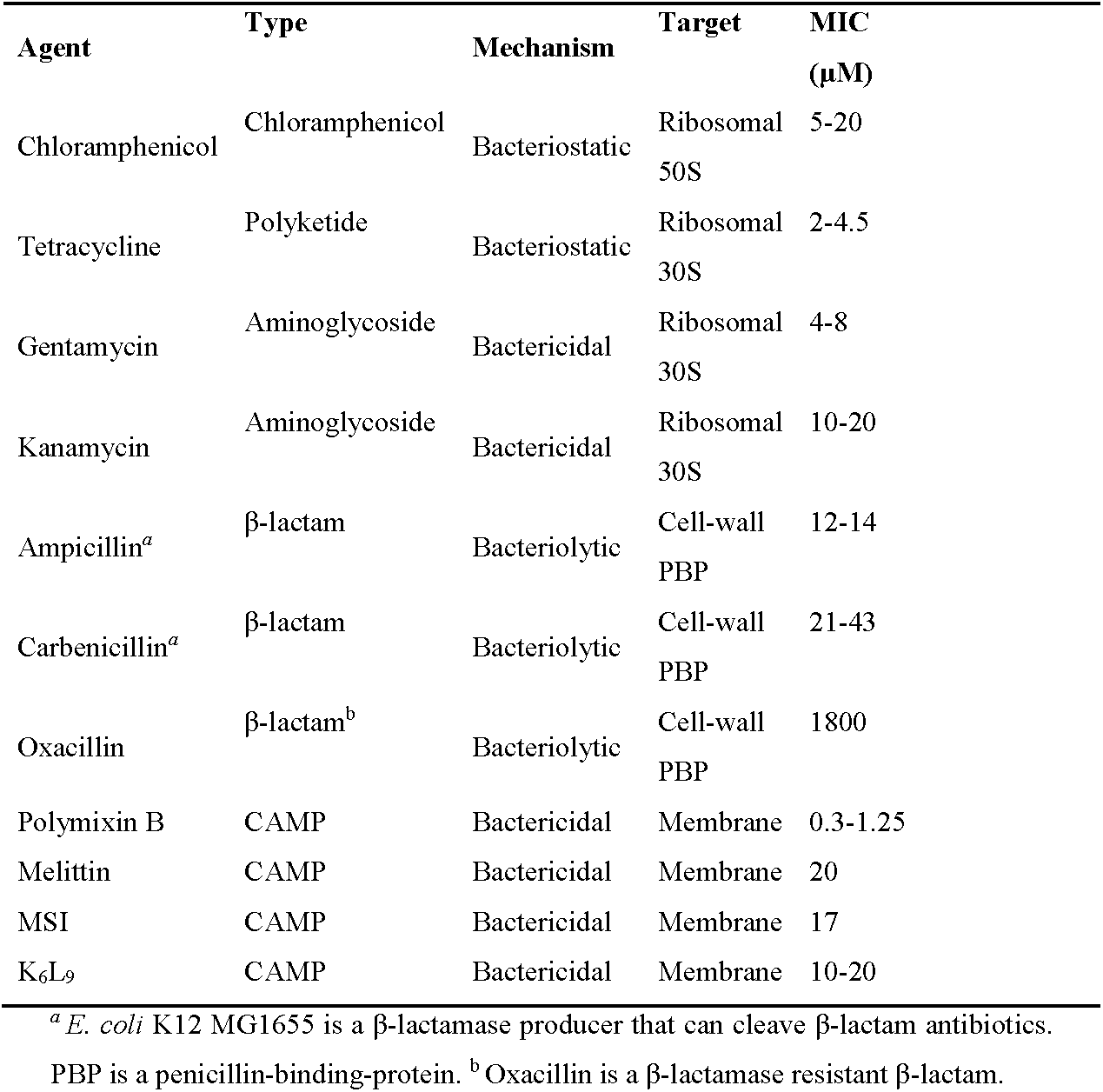
Antibacterial agent properties and tested bMIC ranges concentrations

### Bacterial growth dynamics under antimicrobial treatment

We employed three parameters to characterize the growth dynamics under an antimicrobial treatment (see Flow chart.4):

i. Does the bacterial growth exhibit bi-stability, i.e. is there an antibiotic concentration *A* such that during the experiment time, small initial bacterial loads B(0) do not grow (i.e. are not detectable) whereas large loads do?
ii. Does the bacterial growth exhibit deterministic 1d growth model or more complicated dynamics?
iii. When growth occurs, how does the maximal bacterial concentration depend on B(0) and A?

**Flow chart 4.**
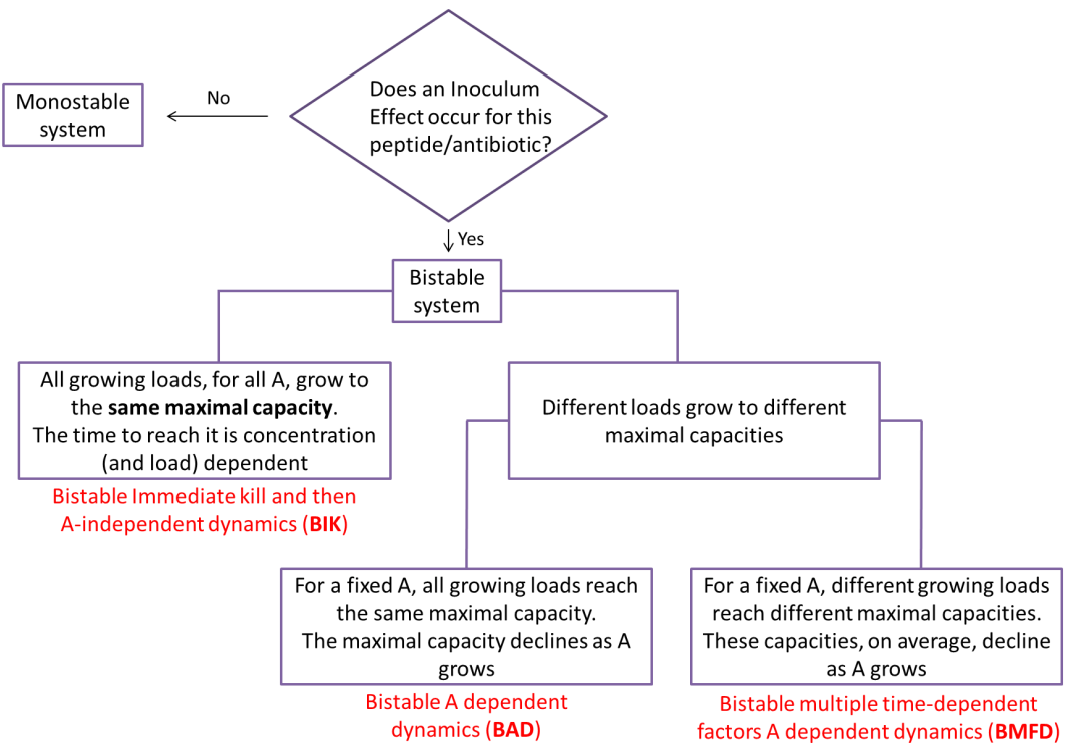
Bacterial growth dynamics in the presence of antimicrobials.

### Bacteria grown in the presence of commercial antibiotics exhibit bi-stability and either A-dependent dynamics (BAD) or more complicated dynamics (BMFD)

In all the experiments with commercial antibiotics bi-stability was detected. Its detection required searching for the bi-stable range which may explain why it was missed in previous studies [22, 33, 34]. Figures 3–4 demonstrate the bi-stable behaviour for Tetracycline and Kanamycin. Similar bi-stability behaviour is observed for all seven commercial antibiotics (see figure S4-S6).

**Figure 3.**
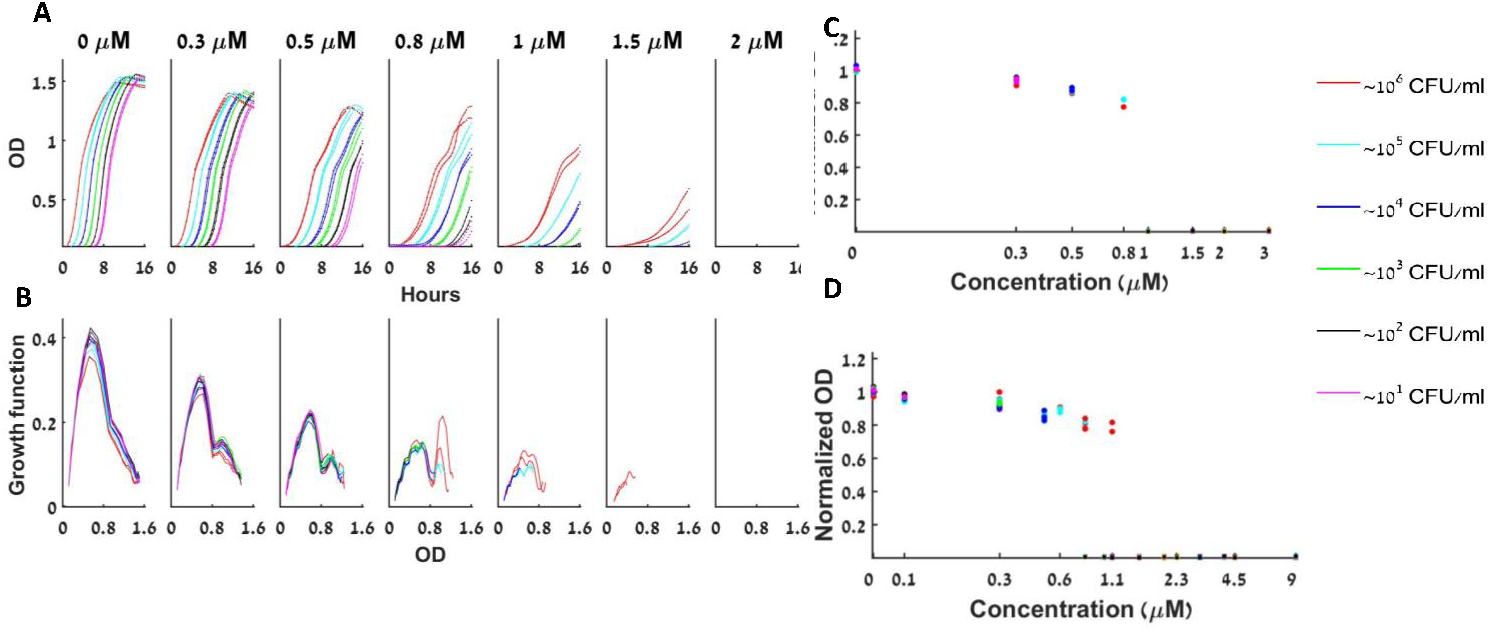
Dynamics of bacterial growth with Tetracycline. (**A**) The bacterial population is either extinct or growing with a similar growth curve under a given concentration of Tetracycline. At any given antibiotic concentration A (one of the seven subplots), all loads that grow have similar growth curves, shifted in time (different colours correspond to different initial loads as indicated). For A>1μm some of the small loads fail to grow altogether. The maximal growth capacity declines as the antibiotic concentration A rises. (**B**) The specific growth function (dOD/dt vs OD) of the different loads changes as a function of *A*, but remains fairly constant for all loads at a given *A*. (**C**) The maximal capacity of all loads at a given Tetracycline concentration A is always similar, and declines as A becomes bigger. Normalized maximal capacities (hereafter, normalized by the averaged maximal capacity of the control) of one representative experiment are presented. (**D**) The normalized maximal capacity of all loads at a given Tetracycline concentration A is always similar, and declines as A becomes bigger – 6 experiments together. *Hereafter, we note that whenever collective results of maximal capacities are represented, all the represented experiments “hit” the dynamic range of the antimicrobial and displayed bistability. See separate experimental results in figure S4.

**Figure 4.**
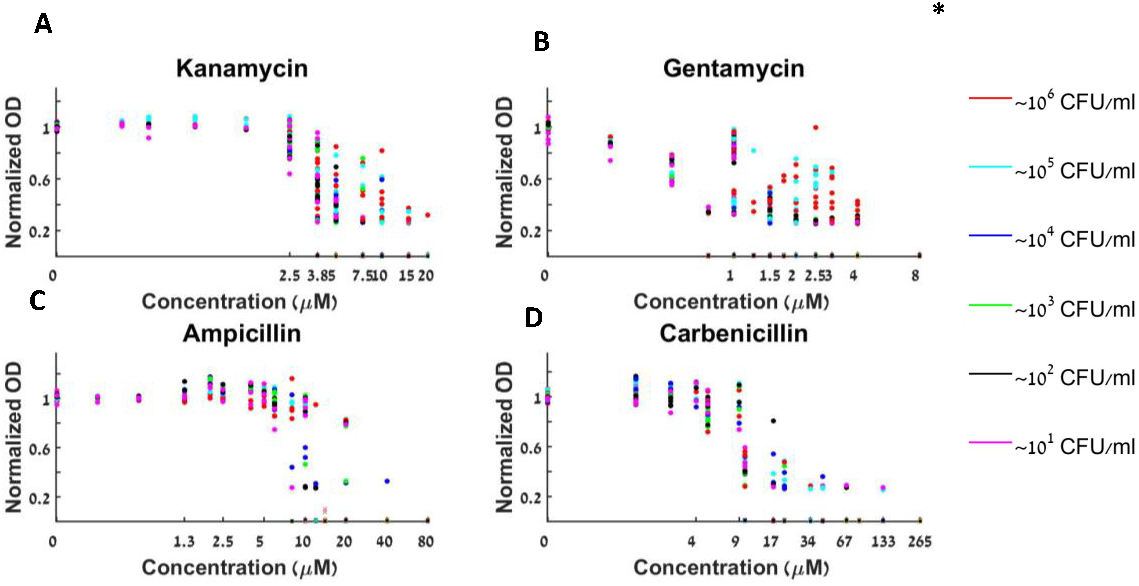
Bacteria grown with β-lactam or aminoglycoside antibiotics exhibit complicated bi-stable dynamics. Collective results of maximal capacity are shown for (**A**) 6 repeats with Kanamycin. (**B**) 4 repeats with Gentamycin. (**C**) 4 repeats with Ampicillin. (**D**) 4 repeats with Carbenicillin. The maximal capacities are load and dose-dependent. * See separate experimental results in Fig S5.

When growing the different initial loads with increasing concentrations of Tetracycline, a bacteriostatic antibiotic that inhibits protein synthesis, we found a bistable behaviour at 1 ≤ A ≤ 3 µM (Figure 3A). Here, for a given *A*, all loads grow with a similar growth function as shown in Figure 3B. Since, for any fixed A, the growth function is not changing when different loads are tested, we conclude that the antibiotic concentration remains roughly constant with time. This behaviour fits our second model – a bistable A dependent model (BAD) in which the growth function is determined only by the antibiotic concentration and is independent of the initial load: dB/dt=F(B;A) where F(B;A) is shown in Figure 3B for several A values. Put differently, for each fixed A value, all the loads that grow, grow in exactly the same way. In particular, their maximal capacity is independent of the initial bacterial load (Fig 3C,D). Moreover, as *A* is increased, the growth rates decline, the smallest loads fail to grow, and the maximal capacity decreases. Repeating the experiment with Chloramphenicol (another bacteriostatic agent) resulted in a similar trend, but at different concentration ranges (Figure S4).

Bistable behaviour was also found using aminoglycoside bactericidal antibiotics Gentamycin and Kanamycin. These agents bind to the 30S subunit of the bacterial ribosome and inhibit translation, which eventually causes cell death. The dynamic range of Kanamycin as deduced from the maximal growth rate is (3.75 ≤ A ≤ 15 µM) (Figure 4A). However, unlike the antibiotics described above, the different initial loads do not grow to the same maximal concentration at a given A. Moreover, the specific growth rates of the different initial loads at a given A do not overlap (Figure 4A). A similar behaviour is demonstrated for Gentamycin with a dynamic range of 1.5 ≤ A ≤ 4 µM (Figure 4B). In this situation, growth dynamics depends on both the initial bacterial load and on other time-dependent factors. Such growth dynamics must be described by at least a two-dimensional growth-inhibition model (or, possibly, by a time dependent one-dimensional model).

The β-lactam antibiotics used in this study (prevent synthesis of new cell-wall peptidoglycan and consequently cause cell-lysis - bacteriolytic), also exhibited a bistable behaviour (see Figure 4C, 4D). With Ampicillin, the bistable range in which large loads grow and small loads go extinct is 8 ≤ A ≤ 12 µM (Figure 4C). Carbenicillin, another β-lactam, is active already at a lower concentration, but has a wider bistable range (8 ≤ A ≤ 16 µM) (Figure 4D). As can be seen from the maximal growth capacity of each initial load at a given antibiotic concentration A, different loads reach different maximal growth capacities (see growth curves in S5), demonstrating a shift from 1d dynamics in the lower antibiotic concentrations towards at least 2d dynamics in the high concentrations. Interestingly, the transition seems to occur in concert with the appearance of the bistability. This kind of dynamics may appear in bistable multiple time dependent factors A-dependent dynamics models (BMFD).

Therefore, bistable behaviour is independent of the specific mechanism of action of each antibiotic. The only condition for such an effect is a suitable range of *A*, which is specific for each agent.

Summarizing, the BMFD dynamics is associated with additional time-dependent factors that influence the bacterial growth. The rich medium we use guarantees that nutrients are not the limiting factor (Fig 1). Naturally, the next factor to test is the antibiotics stability during the course of the experiment (Fig 5A). We find that after 24 hours, Ampicilin potency decreases (MIC doubles) due to exposure to high bacterial loads whereas Carbenicillin potency is reduced due to its exposure to the medium (Fig 5A). Comparing to the stability of the BAD antibiotics (Fig 3c,d), we find that the Chloramphenicol is absolutely stable for at least 24 hours, whereas Tetracycline is less effective after 24 hours (independently of bacterial presence).

**Figure 5.**
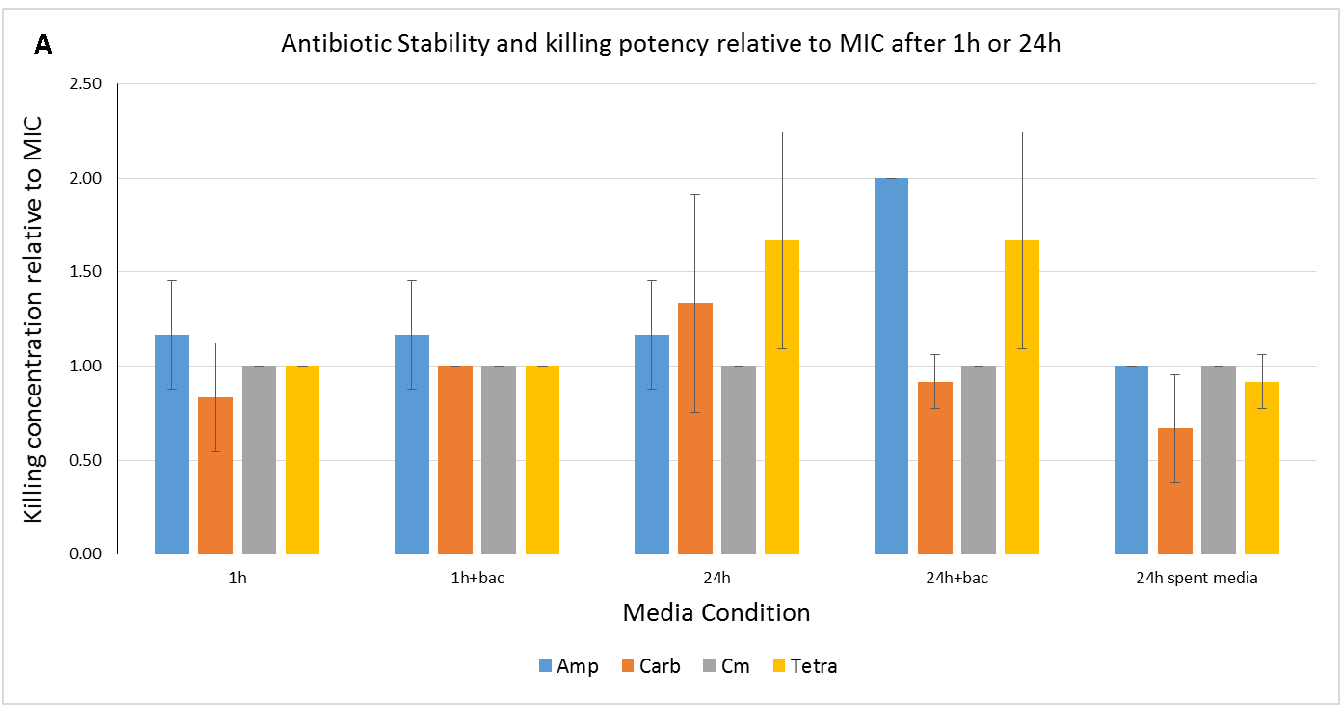
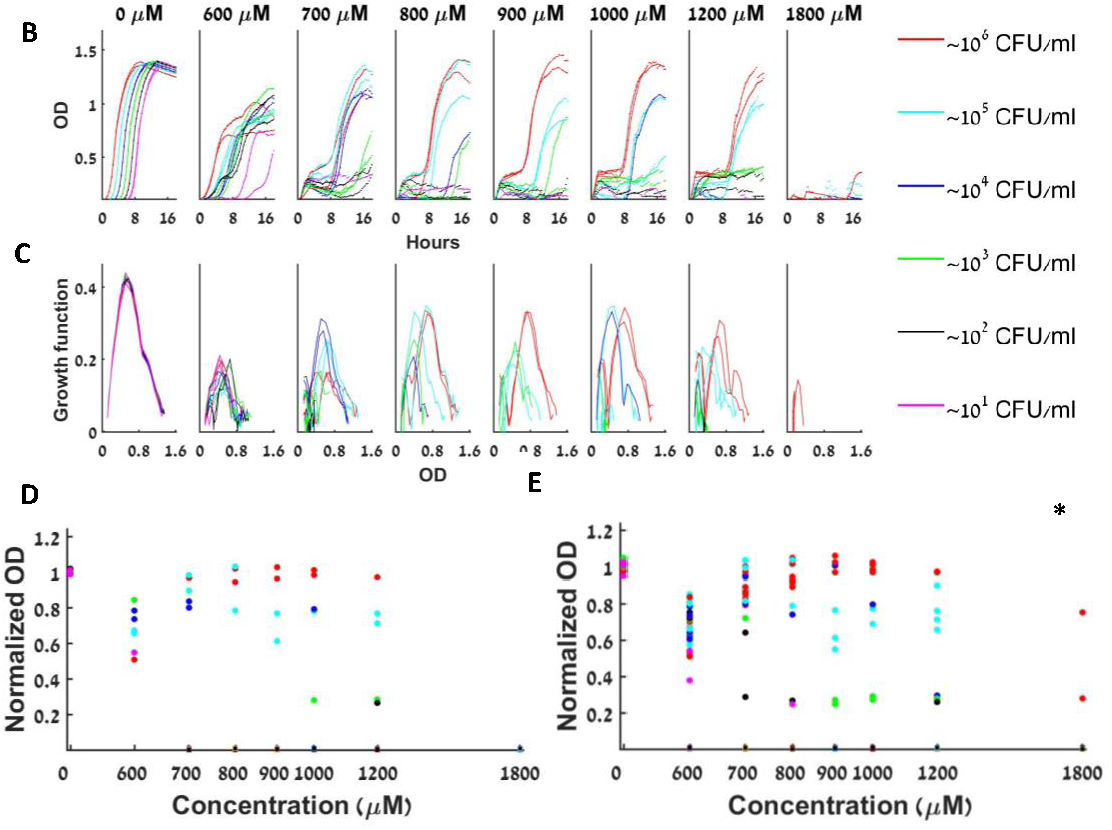
Antibiotic stability over time differs between different classes of antibiotics. (**A**) Minimal inhibitory concentration of the antibiotic relative to the MIC as a function of several media conditions (incubation with or without antibiotics and bacteria as indicated). (**B**) Different loads of bacteria grown with Oxacillin have a similar growth slope in the exponential growth phase at almost all A’s (except for loads that fail to grow altogether). (**C**) The specific growth function of the different loads in Oxacillin changes with the loads and with A, with no obvious trend with respect to A. In particular, the peak of the function remains essentially constant at different Oxacillin concentrations. (**D**) The maximal capacity at all Oxacillin concentrations is always similar, and doesn’t decline as A becomes bigger, although different loads reach different maximal capacities in each antibiotic concentration. (**E**) The maximal capacities of different loads grown in Oxacillin from 6 different experiments. *See separate experimental results in Fig S6.

On the other hand, the stability results also show that substances secreted by the affected bacteria influence the dynamics: the MIC of Carbenicillin is actually reduced in spent medium (Figure 5A right most bar), meaning some bacterial secretions promote the potency of this antibiotic. Also, MIC experiments with Oxacillin, which is a beta-lactam antibiotic resistant to beta-lactamase degradation, demonstrates that even stable antibiotics can produce BMFD dynamics (Figure 5B-E). We conclude that while in some cases the antibiotics potency is reduced during the experiments, additional factors secreted by the affected bacteria may be more significant in producing the BMFD dynamics.

### E.coli K12 grown in the presence of antimicrobial peptides exhibit bistable kill and A-independent dynamics (BIK)

Cationic antimicrobial peptides (CAMPs) directly kill bacteria by perforating or completely disintegrating their membrane [35]. We first tested Polymixin B, a cyclic peptide derived from the bacterium *Bacillus polymyxa* used in the clinic to fight resistant Gram-negative infections. Again, we observed that while large initial bacterial loads could overcome higher concentrations of the peptide, small initial loads could not, even though the exact same amount of peptide was applied (Figure 6A). This implies that antimicrobial peptides also induce a bistable inoculum effect, as do antibiotics. Additionally, all loads at each peptide concentration exhibited the same growth function at each growth phase (see dOD/dt curves in Figure 6B), similarly to the antibiotics that conferred BAD dynamics. Surprisingly, we noticed that the Polymixin B not only induced an inoculum effect, but also did so without changing the maximal capacity of the bacterial growth in all applied concentrations (including concentrations that killed almost all of the initial bacterial loads). Thus the similar look of all dB/dt curves in the presence of Polymixin, except for concentrations of Polymixin B that are 1.25 µM or higher, in which none of the initial bacterial loads survived. Therefore, in the case of Polymixin B, The dynamic range is (0.16 ≤ A ≤ 0.625 µM) where Ae is between 0.625-1.25 µM and Ac is at most 0. 16µM (Figures 6A–C).

**Figure 6.**
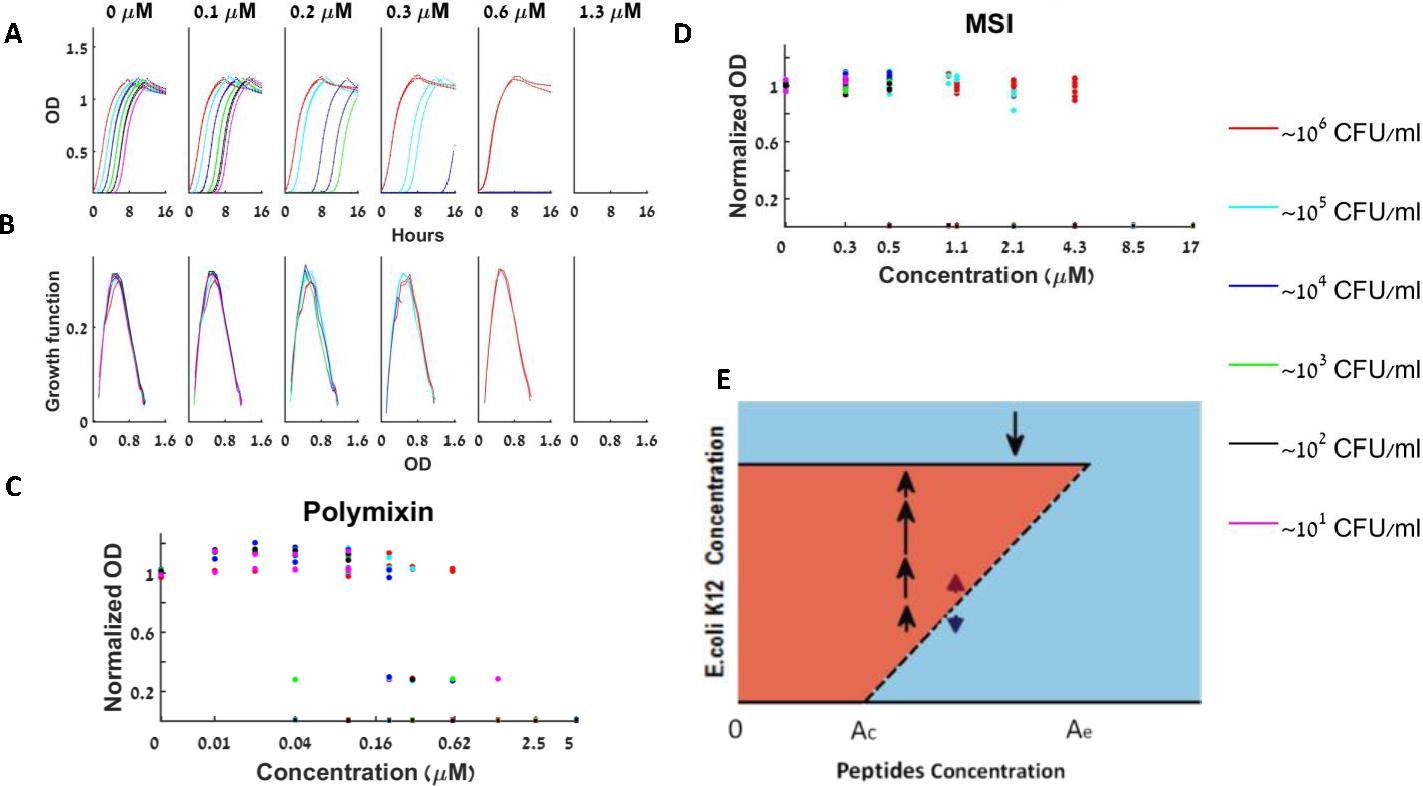
Bacterial growth dynamics under antimicrobial peptides treatment. (**A**) At the bistable range, a given bacterial load is either extinct or grows to full capacity as shown for Polymixin B (**B**) The specific growth functions appear to be independent of both the loads and the peptide concentration (**C**) The normalized maximal growth capacity of each growing load is independent of the Polymixin B concentration – 3 repeats together (**D**) Normalized maximal growth capacities for all loads and peptide concentrations for MSI show similar trend – 4 repeats together. (**E**) The peptide-bacteria interaction can be described by a bifurcation diagram in which bistable behaviour occurs between Ac and Ae. *See separate experimental results in Figure S7.

Interestingly, these same special dynamics were observed for all three other tested antimicrobial peptides – K6L9, MSI and Melittin (Figure 4D and supplementary figure S2) only with different dynamic ranges. This sort of behaviour fits a simplified BAD model. By this model there are still two possible stable bacterial concentrations in the bistable range of peptide concentrations, but these bacterial concentrations are independent of A – bacteria are either completely extinct, or are fully grown by the same growth function as the control, to the maximal capacity of the control well. Namely, when the bacteria grow, they reach a bacterial concentration that is the same for all peptide concentrations. We name this the bistable immediate kill and A-independent growth model, or a model displaying shock dynamics (Figure 4G). Once again, the continuous lines represent stable points for maximal bacterial concentrations, while the interrupted line represents the critical points that determine which of the two possible stable concentrations the bacterial population will take.

This result may be also phrased in terms of the CAMP stability. Since the CAMP loses much of its potency within an hour (Fig S2), the bacteria which survive the first hour grow in the same way as the control.

## Discussion

In all agents we tested, including all cationic antimicrobial peptides and all conventional antibiotics, independently of their biochemical mechanism of action, an “inoculum effect” was found. At a certain range of concentrations, which is specific for every drug and experimental setting, the system exhibits a bistable behaviour in which large loads survive and small loads are inhibited. Moreover, we find that for certain cases (the cationic antimicrobial peptides and the commercial bacteriostatic antibiotics we tested) this phenomenon can be explained by a very simple mathematical model.

Cationic antimicrobial peptides in rich medium (meaning the feeding conditions are not limiting) induce the simplest bi-stable bacterial growth dynamics. Because antimicrobial peptides kill bacteria in a mechanical fashion by attaching to the bacterial membrane via electrostatic forces and formation of pores/lesions through which the bacterial contents flow out, they cause almost immediate death [35]. The only thing that determines the bacterial growth dynamics is the initial peptide concentration “met” by the bacterial cells. After the initial killing is performed by the peptide, there are two possible options: either all bacteria are eradicated, and therefore there would be no growth at all, or, if there are live bacteria left, they will grow normally as the control, independently of the antimicrobial peptide that was primarily present in the medium (see our control experiments in Figure S2). The initial “decision” of whether to go extinct or survive the peptide is highly dependent on the initial bacterial concentration present in the well, and therefore, an inoculum effect occurs. Since the initial load is reduced abruptly by the killing, the time till the bacterial load becomes detectable allows an estimate of the killing rate – this rate appears to be highly sensitive to the experimental setting (see Supplementary figure S7). Mechanistically, this model is supported by various biophysical studies done by others and ourselves showing that CAMPs bind and kill both Gram-negative and positive bacteria within minutes [36, 37] see Figure S2. The consequence of this unusual interaction is mathematically described by our uni-dimensional BIK model – there is a density dependent and concentration dependent immediate kill function, with the surviving bacteria growing exactly as the control, obeying a deterministic one-dimensional ordinary differential equation with a growth function that is found experimentally (Figure 6B).

Commercial bacteriostatic antibiotics that target the ribosome also induce a simple bistable bacterial growth dynamics - a concentration dependent unidimensional growth-inhibition activity. Such behaviour appears when there is no decline in the effective concentration of the antibiotic in the medium and the medium is rich. Then, the bacterial growth function depends only on the instantaneous inoculum and on the amount of antibiotic available at the time of exposure, [A_0_]: at any given [A_0_]different initial bacterial loads obey the same dynamic laws which can result in inhibition or growth to an [A_0_] dependent maximal concentration. Such behaviour is described mathematically by our BAD model - deterministic one-dimensional ordinary differential equation with growth function that depends on the initial antibiotics concentration [A0]. Notably, we find these functions from the experimental data, and observe that they may have multiple maxima for large [A0] (Figure 3 and S4). These experimentally derived functions may be utilized in future mathematical models of bacterial growth dynamics.

Treatment with beta-lactam antibiotics, which specifically target the cell-wall, or with aminoglycosides that inhibit the ribosome, resulted in a more complex bi-stable growth dynamic. Our results demonstrate that the growth function of the bacteria for any given initial drug concentration [A_0_] does not depend only on the time-dependent bacterial density, namely it cannot be described by a single time independent ordinary differential equation. Such a behaviour may be explained by more complex mathematical models, which we call BMFD. Such models include additional time-dependent factors, for example, the time-dependent antibiotic concentration (see, e.g. [7, 12] for a variety of such models).

A summary of the three described models and the division of the tested antimicrobial agents between them is visualized in Flow Chart No 5.

**Flow Chart No 5.**
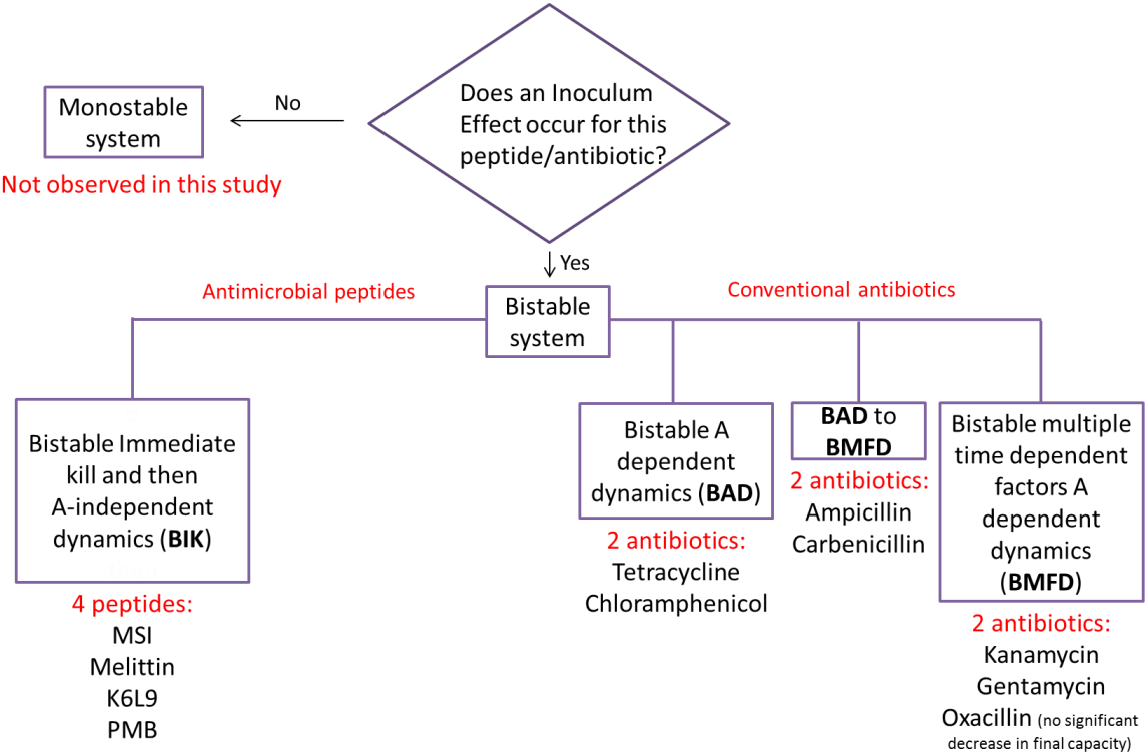
Summary.

While IE was identified in previous studies of particular conventional antibiotic agents and bacteria [6, 7] [9, 10] [7, 8, 12], previous explanations of its appearance included genetic and/or phenotypic population heterogeneity and additional time-dependent factors. These were modelled, for example, by deterministic multi-dimensional equations of classical reaction kinetics [7, 12] or by two-dimensional PK/PD dynamics [8]. Here we show that for some cases (the bacteriostatic antibiotics) the one dimensional BAD model can explain the resulting growth curves by density dependant mechanisms alone. By Ockham’s razor principle, we assert that the BAD models are adequate for these cases: the more complex reasoning can be neglected for explaining bacterial growth in rich media under bacteriostatic antibiotics.

On the other hand, we demonstrated that in other cases (bi-stable bacterial growth with bactericidal and bacteriolytic antibiotics), higher dimensional models are required. These could possibly be described by either density dependant mechanisms (such as the production of β-lactamase or, for other bacteria, by extracelluar PH variations[8]) or models involving population heterogeneity, see [7, 8, 12] and references therein. For example, it is well established that *E. coli* MG1655 produces β-lactamase, an enzyme that hydrolyses β-lactam antibiotics such as Ampicillin and Carbenicillin [7, 32]. If the antibiotic is being inactivated by the bacteria, higher initial loads would then produce larger concentrations of the enzyme over the course of time, and therefore, the antibiotic concentration will decline faster. Our experimental results with Oxacillin demonstrate that even when the β-lactam is not degraded by the bacteria and the drug concentration remains constant over time, IE appears, and the growth dynamic may be still multi-dimensional. Interestingly, these results also exhibit high sensitivity to the experimental settings. This sensitivity could be possibly attributed to the stress induced by the very high antibiotic concentrations (*E.coli* are often naturally resistant to Oxacillin [38]), possibly introducing an additional time and antibiotic concentration dependant factor to the system. Additional experimental work in which these factors are monitored is needed for clarifying the dominant mechanisms involved in the more complex settings.

The BAD models and their extended multi-dimensional BMFD models suggest a basic mechanism of bacteria-antibiotics interactions, or for that matter, the dynamics of bacterial growth under any kind of limiting conditions and not just antibiotics (for example, the BAD dynamics are similar to in-vitro neutrophils-bacteria interactions [30] and the BMFD may be relevant to our experiments with minimal medium (Figure 2C-D).

From a therapeutic point of view, understanding the particular bacterial growth dynamic in the presence of different classes of antimicrobials and starting from various bacterial inoculums is central for the assignment of the correct treatment for various bacterial infections (see [7, 12] and references therein). Of-course, bacterial infections *in-vivo* require additional considerations, such as the presence of the immune system, the clearing of the drugs, as well as environmental conditions that are not only subjected to changes by the bacteria but also by the host (e.g., the host may control limiting factors for the bacterial growth, and such factors invoke higher dimensional dynamics, see Figure 2C-D). Nevertheless, studies have shown [14, 39] that quite often *in-vitro* results regarding interaction dynamics between bacteria and antimicrobials are indicative of such dynamics *in-vivo* and may be used as building blocks for the in-vivo models [30]. If the IE is relevant for application of any antimicrobial agent *in-vivo* as we show here is the case *in-vitro*, then the simplest way to avoid its alarming consequences is to treat infections when they are still very small. This would reduce the chance of failure in standard clinical treatment protocols and will not allow bacteria to acquire genetic resistance over a long term of insufficient treatment. However, in cases when early diagnosis and treatment are impossible, the ability to predict bacterial growth dynamics in the presence of a selected treatment becomes indispensable. In such cases, the estimation of bacterial load present in the infection site, and the knowledge of the type of growth dynamic of the infecting bacterium with different antimicrobials, would allow for a personalized treatment in terms of dosage and frequency of treatment [22, 40] [6, 7]. Examples for the possible growth models include BIK, BAD or BMFD dynamics.

Our current efforts are concentrated on simulating the results based on the fitting of our experimental data and studying their implications. Interestingly, in many of the antibiotic concentration dependent growth functions a “hump” in the graph is seen at higher OD’s (see Figure 3). This can be indicative of a change in the collective behaviour of the bacteria at these higher densities. Investigation of this change in the growth dynamic in the future might shed light on the distinct bacterial behaviour at high densities. Finally, to achieve a successful clinical treatment, basic growth dynamics rules such as those presented herein should be adapted to include additional parameters such as nutrient limiting factors, drug clearance, the action of the immune system and the level of drug resistance of the specific bacterial species present at the infection site. Inclusion of these additional factors into the bi-stability of the basic bacteria-antimicrobial agents system can shed light on the relation between in-vitro and in-vivo growth dynamics.

## Acknowledgement

We thank the SELA-YEDA Centre at the Weizmann Institute of Science for their support. We thank Roy Malka for discussions and comments.

